# Phenotyping ciliary dynamics and coordination in response to CFTR-modulators and Thymosin-α1 in Cystic Fibrosis respiratory epithelial cells

**DOI:** 10.1101/223024

**Authors:** M. Chioccioli, L. Feriani, J. Kotar, P. E. Bratcher, P. Cicuta

**Affiliations:** Biological and Soft Systems Sector, Cavendish Laboratory, University of Cambridge, Cambridge CB3 0HE, UK; Division of Cell Biology, National Jewish Health, Denver, CO, USA

**Keywords:** Cystic Fibrosis, Cilia, VX809, VX770, Tα1, CBF, cilia coordination, ALI, multi-DDM

## Abstract

The diagnosis and treatment of respiratory disorders are challenging and would benefit from new approaches to systematically assess ciliary beating dynamics and to test new drugs. A novel approach based on multiscale differential dynamic microscopy (multi-DDM) is shown to quantitatively assess collective beating of cilia in a non-biased automated manner, in human airway epithelial cells (HAECs) derived from subjects with cystic fibrosis (CF) and grown in 2D air-liquid interface culture. Multi-DDM can readily detect changes in both ciliary beat frequency (CBF) and cilia coordination that result from perturbations to the mucosal layer. The efficacy of three CFTR-modulating treatments is investigated: ivacaftor (VX-770) with lumacaftor (VX-809), VX-809 alone and Thymosin alpha 1 (Tα1) alone. All three treatments restore coordination of cilia beating in the CF cells, albeit to varying degrees. We argue cilia are affected by these treatments through the physical properties of the mucus. Phenotyping cilia dynamics through multi-DDM provides novel insight into the response of ciliary beating following treatment with drugs, and has application in the broader context of respiratory disease and for drug screening.

**One sentence summary:** A semi-automated and unbiased assay based on multiscale differential dynamic microscopy (multi-DDM) detects changes in the coordination and frequency of ciliary beating in F508del/F508del primary human airway cells under different conditions and in response to CFTR-modulating compounds.

## Introduction

Respiratory disorders affect millions of people worldwide and can result from both genetic and environmental causes *(1, 2)*. Many respiratory disorders are characterised by abnormal ciliary beating, be this a causal or derivative behaviour. Current approaches to systematically analysing collective cilia beating are unfortunately limited, as is our understanding of the physical properties required to sustain healthy transport of mucus out of the airways, known as mucociliary clearance (MCC)*(3)*. Most approaches to phenotyping ciliary motion and coordination are time-consuming and difficult to standardise across labs *(4)*. Videomicroscopy examination of airway biopsies is often used to estimate ciliary beat frequency (CBF) and to inspect waveforms on individual cilia, but this is usually performed manually and requires experienced personnel. Semi-automated approaches to measure CBF have been developed recently, however even these assess only a subset of the sample and thus cannot detect the broad distribution of CBF that can occur within a given biopsy *(10, 14 - 17)*. Furthermore, while CBF is a first and readily accessible phenotype, it is not sufficient to diagnose pathologies: it is clearance as a whole that has to function.

A key parameter of healthy mucociliary clearance relates to how the ciliary beating is coordinated across large (many cell) distances. Despite its importance, the characterisation of cilia coordination in the context of human respiratory disease has not been well explored. Approaches probing both CBF and cilia coordination have been reported, however these either require manual selection of the area for analysis *(4)*, or are not suitable for samples grown in 2D air-liquid interphase (ALI) culture, which is the standard method for culturing clinical HAEC samples *(10–12)*. In ALI culture, the ciliated cells typically exist in patches, and coordination across the entire sample is highly variable (*13, 14*), precluding for example assays based around bead-clearance. As such, there is very little data to describe how collective and coordinated ciliary beating arises in healthy human airway cells, and how this goes awry in disease. We recently reported a video analysis algorithm based on differential dynamic microscopy (DDM), which we called multiscale DDM (multi-DDM)*(15)*. This allows the characterization of collective ciliary beating in human airway epithelial cells in a fast and fully-automated manner. The input required to run the DDM or multi-DDM algorithms are typically 10 second bright-field optical microscopy videos of the ALI cultured cells in situ, taken at 40x magnification and moderately high frame rate (∼150 frames per second). By considering the frame differences at various time intervals, transformed in Fourier space, the method extracts temporal and spatial coherence of any dynamics in the video. In particular, for videos with oscillating features such as motile cilia, the CBF is obtained without the need to neither segment nor select regions. In multi-DDM *(15)* we showed that the spatial scale of coordinated cilia dynamics can be measured.

We focus this study on cystic fibrosis (CF): a life-threatening genetic disorder caused by loss-of-function mutations in the cystic fibrosis transmembrane conductance regulator (*CFTR*) gene *(16)*. Defective CFTR activity in airway epithelial cells leads to loss of airway surface liquid and incompletely hydrated mucin, resulting in a thick layer of mucus that obstructs the airways and promotes chronic bacterial infections and inflammatory lung damage. In individuals with CF, cilia beating is greatly compromised to the extent that the mucus cannot be properly cleared, and the range of cilia movement is severely restricted *(16, 17)*. Of the nearly 2000 CFTR mutations that have been identified (www.genet.sickkids.on.ca), only the most common mutations expressed by large groups of subjects have been targeted for drug screening due to the high cost and time-consuming nature of clinical trials. However, a recent study showed that selected CFTR-modulating drugs used to treat patients with the common F508del-CFTR mutation could in some cases be effective in treating patients with rare, uncharacterized CFTR mutations that are currently not registered for treatment, if only cost-effective assays could be used to screen such samples *(18)*.

In the present study, we performed multi-DDM analysis showing that primary HAECs obtained from subjects with the F508del/F508del mutation in CFTR exhibit unique cilia coordination and CBF dynamics compared to cells from healthy subjects. multi-DDM data quantify in a very direct way the loss of cilia coordination with distance, and how this phenotype is affected by mucus properties in both direct perturbations and pharmacological intervention, providing new important information about coordination of cilia dynamics in the context of CF. As a proof-of-concept, we used this approach to demonstrate the efficacy of the CFTR-modulating drug combination of VX-770 (ivacaftor/ KALYDECO^®^) and VX-809 (lumacaftor, together termed ORKAMBI^®^) on HAECs derived from subjects homozygous for F508del CFTR. We further investigated the effect of VX-809 alone on these cells, as well as a recently-reported CFTR-modulating compound Tα1 *(19)*. The development of a rapid, quantitative assay for characterising collective cilia beating dynamics in a standard cell culture model has important applications in diagnostics and drug screening. To our knowledge, these data represent the first in-depth, quantitative assessment of cilia coordination in HAECs derived from subjects with CF and how this changes in response to CFTR-modulating drugs. Although this study applies multi-DDM analysis to investigate ciliary beating dynamics in the context of CF, our approach to phenotyping cilia dynamics could be applied to the other respiratory diseases in which ciliary beating is affected.

## Results

### Multi-DDM analysis of CBF and cilia coordination in healthy HAECs

One of the first parameters normally probed in clinical samples of HAECs, and one which is a general indication of ciliary function, is the CBF. Unlike standard clinical practice whereby a single CBF value is calculated from a user-selected region-of-interest (ROI) *(20, 21)*, multi-DDM provides the user with a distribution of CBF values measured across the imaged fields of view (FOV). This is important, since CBF can vary across a single sample (Figure 1A – B), and thus a single point measurement is not an accurate representation of the entire sample. Dividing the entire FOV into square subsets of a pre-defined size (called tiles - 64 × 64 pxl, 9.3 µm per side; Fig. 1C) and running the DDM algorithm on each of these tiles generates a distribution of measured CBF across the entire FOV (Fig. 1D, E). In the measurement of CBF, DDM simply extends in a systematic and user-free fashion the concept of probing intensity modulation over time, in a pixel or group of pixels.

**Figure 1.**
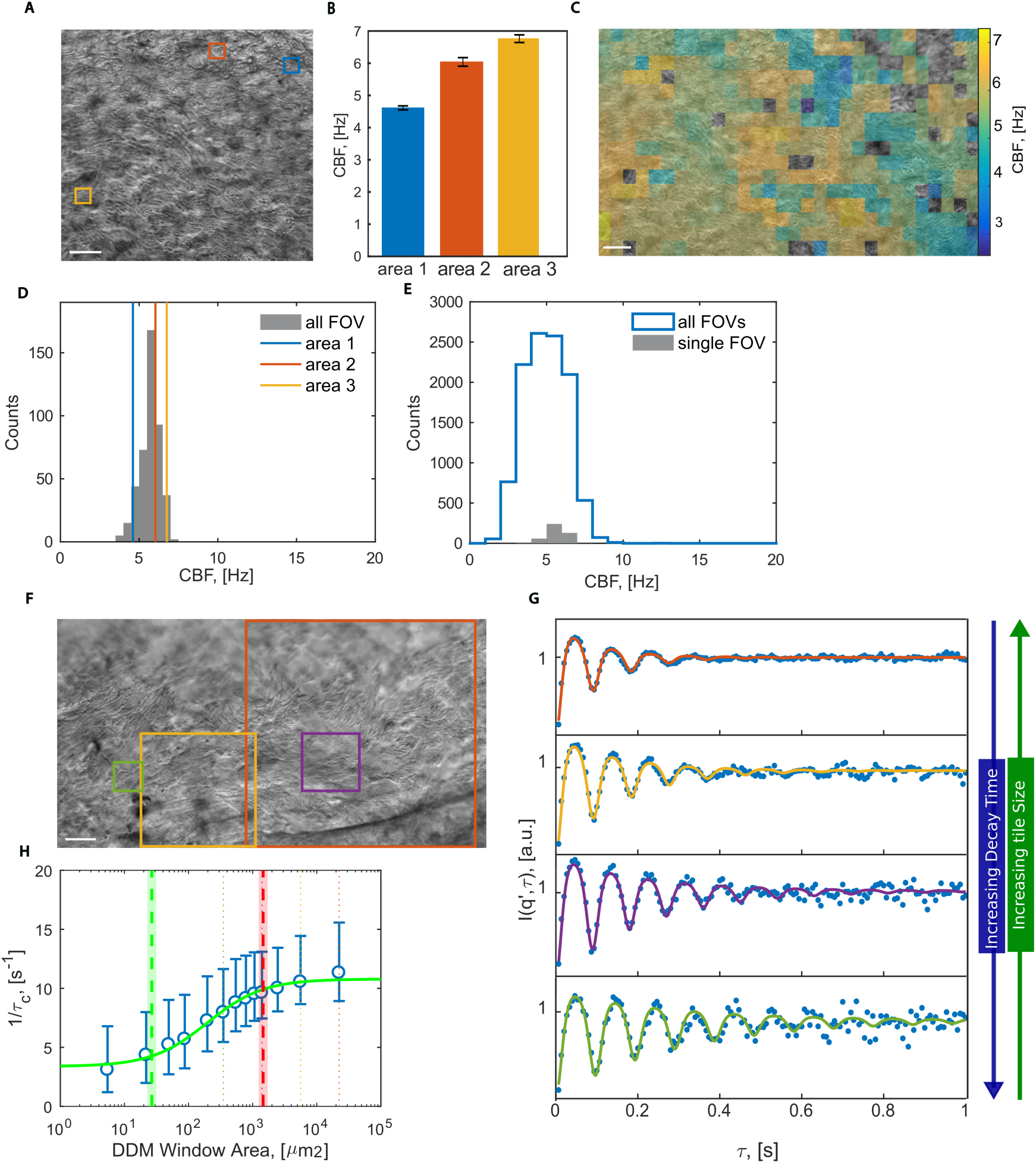
Quantitating ciliary beat dynamics in healthy HAECs via multi-DDM. **(A)** The standard approach for determining CBF is to manually select a region-of-interest (ROI) for analysis from a microscopy video. An example image from a microscopy video used in the analysis with three different 64 x 64 pixel regions (red, blue and yellow) selected for CBF analysis. Scale bar is 20 µm. **(B)** The regions of interest selected for analysis in (A) displayed a range of CBFs: 4.6 ± 0.1 Hz, 6.0 ± 0.1 Hz, and 6.75 ± 0.1 Hz. CBFs differed among the three regions, highlighting the subjective nature of analysing CBF via user-selected regions of interest. To capture the distribution of CBFs in an objective manner, the entire field of view was divided into many square tiles and DDM was performed on each tile to obtain CBF. The division in tiles in a FOV, and the respective measured CBF values (in false colours), are shown in **(C)**. Tiles where no motion was detected remain transparent. The full distribution of CBF values measured from this FOV is shown in (**D**), where the CBF measurements from the 3 tiles of (**A, B**) are shown with vertical lines. The mean of this distribution was 5.7 Hz, with 90% of the values falling between 4.6 Hz and 6.6 Hz. A more exhaustive measurement of CBF over the entire ALI sample can be obtained by taking several microscopy videos from different FOVs scattered around the insert. (**E**) shows the distribution of CBF values built by aggregating the distributions from all such FOVs. We also show on the same axes, for sake of comparison, the same CBF distribution as in (**C**). The aggregated distribution had a mean of 4.9 Hz, with 90% of measured values falling between 2.8 Hz and 7.0 Hz. To measure the coordination length scale of beating cilia we first divided the FOV into tiles, repeating this process while systematically increasing the size of the tiles. DDM is then run on each tile. For example, the FOV in (**F**) shows the outline of 4 tiles, each of a different size. As extensively detailed in *(15)* we can fit the time dependence of the output of the DDM algorithm (Image Structure Function) with an oscillating function multiplied by an exponentially decaying term. The frequency of the oscillations is the CBF, while the time constant of the decaying term is a proxy for how well coordinated the motion within the analysed region is. **(G)** Examples of the oscillating and decaying behaviour of the Image Structure Function for each region shown in **(F)**. Fitted curves (solid lines, colours matching the respective regions in (**F**) were superimposed onto the experimental data (dots), and showed excellent agreement. Note how the time constant increases as the size of the tiles decreases. By plotting the median of the inverse time constant *τ*_*c*_ (circles, while whiskers show the 25^th^ and 75^th^ percentile) measured at each tile area across several fields of view it is possible to build a sigmoidal curve that quantitatively represents the length scale of ciliary coordination **(H)**. The left and right shoulders of the sigmoidal curve are indicated by the green and red dashed lines respectively, while the shaded region indicates the 68% confidence interval. The area of the highlighted tiles in (**F**) is reported here as a dotted line of matching colour.

The standard clinical practice for analysing CBF in ALI cultures does not capture any information regarding the spatial coordination of cilia beating. However it is clear that cilia across the airway epithelium must beat in a coordinated fashion in order to efficiently propel the mucus layer upwards, and uncoordinated ciliary beating is a hallmark of some ciliopathies, for example PCD *(22–24)*. To assess the degree of spatial coordination we deployed multi-DDM, where the size of the tiles on which the algorithm is run is systematically changed (Fig. 1F): as the tile probed is decreased in size, cilia beating within the tile becomes more coordinated, which is seen as an increase in a characteristic decay time of the resulting signal (Fig. 1G). Plotting the inverse of these decay times against their tile size produces a sigmoidal dataset that functions as a quantitative measure of cilia coordination within a given sample (Fig. 1H). This data can be fitted empirically to extract a parameter (essentially the position of the sigmoidal curve) that pinpoints the spatial scale of cilia dynamics coordination. The “shoulders” in the sigmoids are defined as the points where the line that best approximates the central slope meets the respective baselines: the left shoulder (λ^2^; Fig. 1H, green dashed line) marks the square of the length scale at which motion starts losing coordination, whereas the right shoulder (Λ^2^; Fig. 1H, red dashed line) marks the square of the maximum length within which the sample still shows some coordination. These parameters λ and Λ are found by fitting the experimental sigmoids with 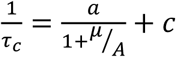, where *a* and *c* give the level of the plateaus, *A* is the area of the DDM tile, and *µ* is the inflection point of the sigmoid. Since λ and Λ are separated by a constant shift on the *A*-axis, it is possible to use either of the two in this work to compare between different experimental conditions, depending on which one falls within the accessible range of our instrumentation.

### Multi-DDM analysis of CBF and cilia coordination in CF HAECs

We used multi-DDM to compare CBF and cilia coordination in HAECs from healthy donors and from subjects with CF. In CF cells defective CFTR activity leads to loss of airway surface liquid and incompletely hydrated mucin, resulting in a thick mucus layer that greatly restricts cilia beating. The distribution of CBFs for a typical sample with the F508del/F508del CFTR mutation is shown in Fig. 2A (bottom left panel). Multi-DDM analysis of HAECs obtained from three subjects with the same mutation reveals average CBFs ranging from 3.94 Hz to 6.28 Hz. By comparison, HAECs obtained from healthy subjects exhibits a higher average range of CBFs, from 5.06 Hz to 6.56 Hz. A typical distribution of CBFs for a healthy HAECs sample is in Fig. 2A (top left panel). Average CBF and standard deviation are reported in Fig. 2B for all subjects, and in Supplementary Table 1 together with 90% range.

**Figure 2.**
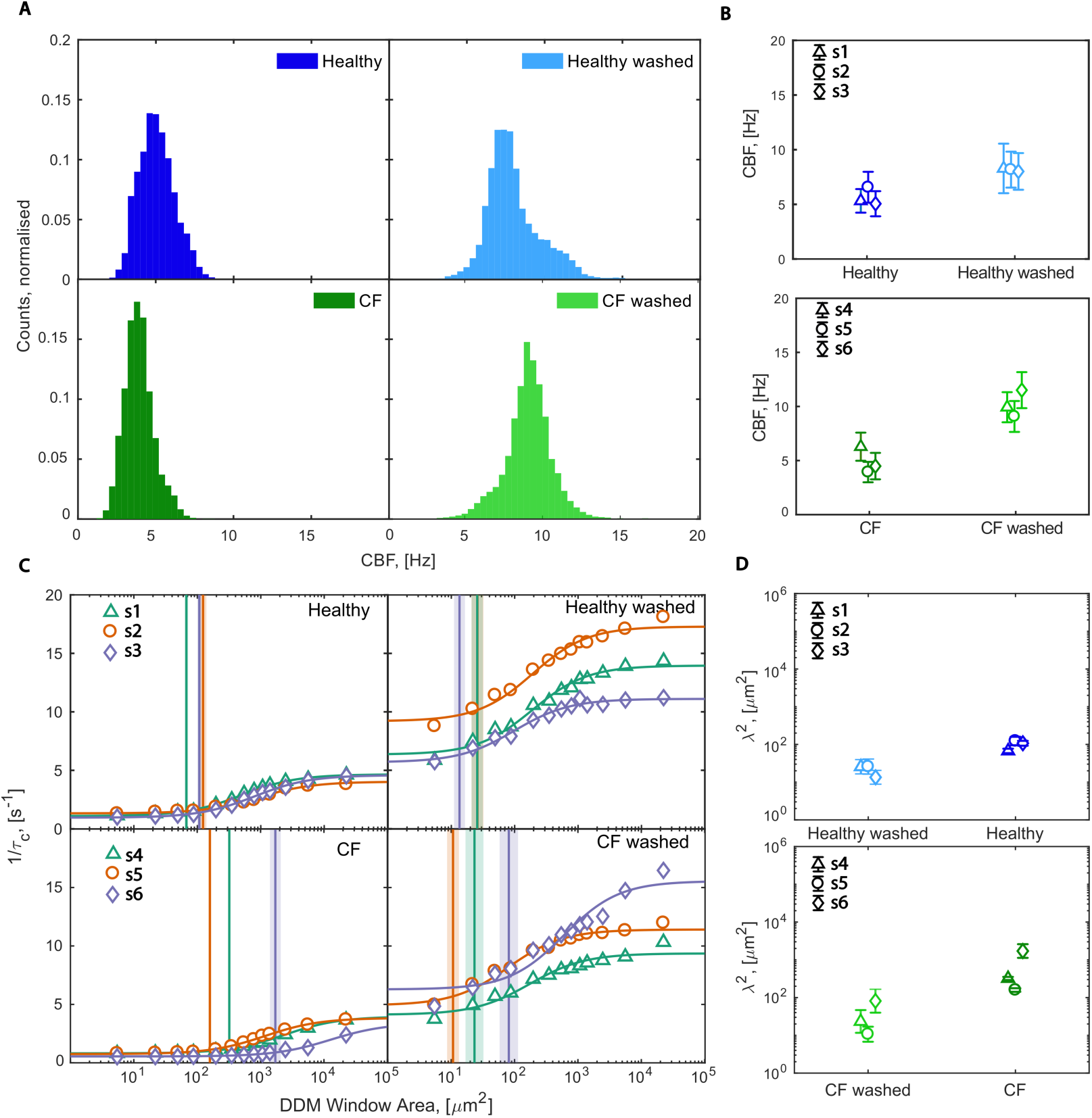
Rheological properties of mucus affect CBF and the length scale of ciliary coordination. **(A)** The distribution of CBF from cells obtained from healthy subjects compared to subjects with CF (F508del/F508del) before and after mucus removal. Each distribution was built from at least 20 fields of view imaged across the respective sample. **(B)** The mean and standard deviation for the CBF measured before and after mucus removal on each of the three separate healthy donors (s1, s2, s3) and CF (s4, s5, s6). **(C)** The sigmoidal curves of the inverse of the time constant *τ*_*c*_ as a function of the area of the DDM tile for samples obtained from three healthy subjects (left) and three subjects with CF (F508del/F508del) (right), before and after mucus removal (top and bottom, respectively). Experimental data are shown as symbols (for the sake of visual clarity, error bars showing 25^th^ and 75^th^ percentiles were omitted), and the sigmoidal fit is shown as a continuous line. The vertical line represents the left shoulder point and marks the coordination length scale squared *λ*^2^, and the shaded area its uncertainty. Each sigmoidal curve is built from at least 20 FOVs imaged across the sample. CF samples are characterised by a higher length scale of ciliary coordination than healthy samples, however this effect is only apparent in the presence of mucus. (**D**) Comparison of the *λ*^2^ for each sample revealed significant changes in the length scaled of ciliary coordination following removal of the mucus (P < 0.05 and P < 0.01 for healthy and CF samples respectively. Paired, two-tailed Student’s t-test comparing log_10_(*λ*^2^).

We hypothesise that the differences in ciliary beating between cells from healthy subjects versus subjects with CF is attributable to the physical properties of the mucus, since the CFTR mutation affects mucosal properties and not the intrinsic structure or function of cilia *(25, 26)*. To test this, we analysed F508del/F508del HAECs both in the presence of mucus and immediately after a wash treatment to remove mucus. Average CBFs obtained following a wash treatment are typically higher than when mucus is present, and this is true for cells obtained from both healthy and CF subjects (Fig. 2B and Supplementary Table 1). We next assessed the degree of ciliary coordination in the CF samples in the presence of mucus and immediately following a wash. The curves generated using the multi-DDM algorithm show the typical sigmoidal shape for both conditions, however we observe that the position of the left shoulder of the sigmoid (λ^2^) differs depending on the presence or absence of mucus (Fig. 2C). In the absence of mucus (i.e., immediately following a wash) the ciliary coordination length scale decreases towards values obtained from healthy samples. Direct comparison of λ^2^ between the two conditions reveals a statistically significant (p<0.01) shift to smaller values when the mucus is removed in the HAECs obtained from CF subjects. This is also observed to a lesser extent when mucus is removed from HAECs obtained from healthy subjects (p<0.05; Fig. 2D). The shift to smaller values represents a decrease in the length-scale of ciliary coordination, meaning decreased coordination in the absence of mucus. This shift implies that the mucus, acting like an elastic gel raft, helps cilia to synchronise their dynamics. The difference between the shoulder values obtained from two treatments of a given sample can be used to quantitatively differentiate between ciliary beating dynamics.

### Multi-DDM charts the evolution of ciliary beating phenotypes over time

The experiments with multi-DDM presented above analysed and compared different HAEC samples maintained under constant cell culture conditions, which is useful to obtain a snapshot comparison between two samples but does not convey information about how the cilia dynamics phenotype evolves over time in response to various conditions. This is particularly relevant for understanding cellular responses to pathogenic infections and/or pharmacological responses. Having identified that the presence of mucus has a dramatic effect on both CBF and cilia coordination, we tested the effect of successive cycles of washing and allowing the mucus to regenerate. HAEC samples obtained from three different subjects with the F508del/F508del mutation were subjected to successive rounds of mucus removal and regeneration for two days. Briefly, DDM data were collected in the presence of mucus (0 hrs) and then immediately after mucus was removed (+2hrs), then again once the mucus had regenerated (+24hrs). This cycle was repeated at 26 and 48hrs (Fig. 3A). We observed complete regeneration of the mucus (consequently compromised cilia beating) within 24hrs, regardless of whether this occurred on the first, second or third wash. The DDM analysis provides a quantitative measure of this, revealing cyclical changes in CBF over time, with the average CBFs ranging from 2.4 – 5.1 Hz in the presence of mucus, and 8.4 – 10 Hz once mucus was removed (Fig. 3B, D and Supplementary Table 2), consistent with our previous observations. There is no significant difference between the change in CBFs over successive cycles of mucus washing and regeneration, nor between different donor samples (Fig. 3C, D), indicating both the sensitivity (consistent over different time points) and the robustness (consistent across different donors) of the DDM approach over time.

**Figure 3.**
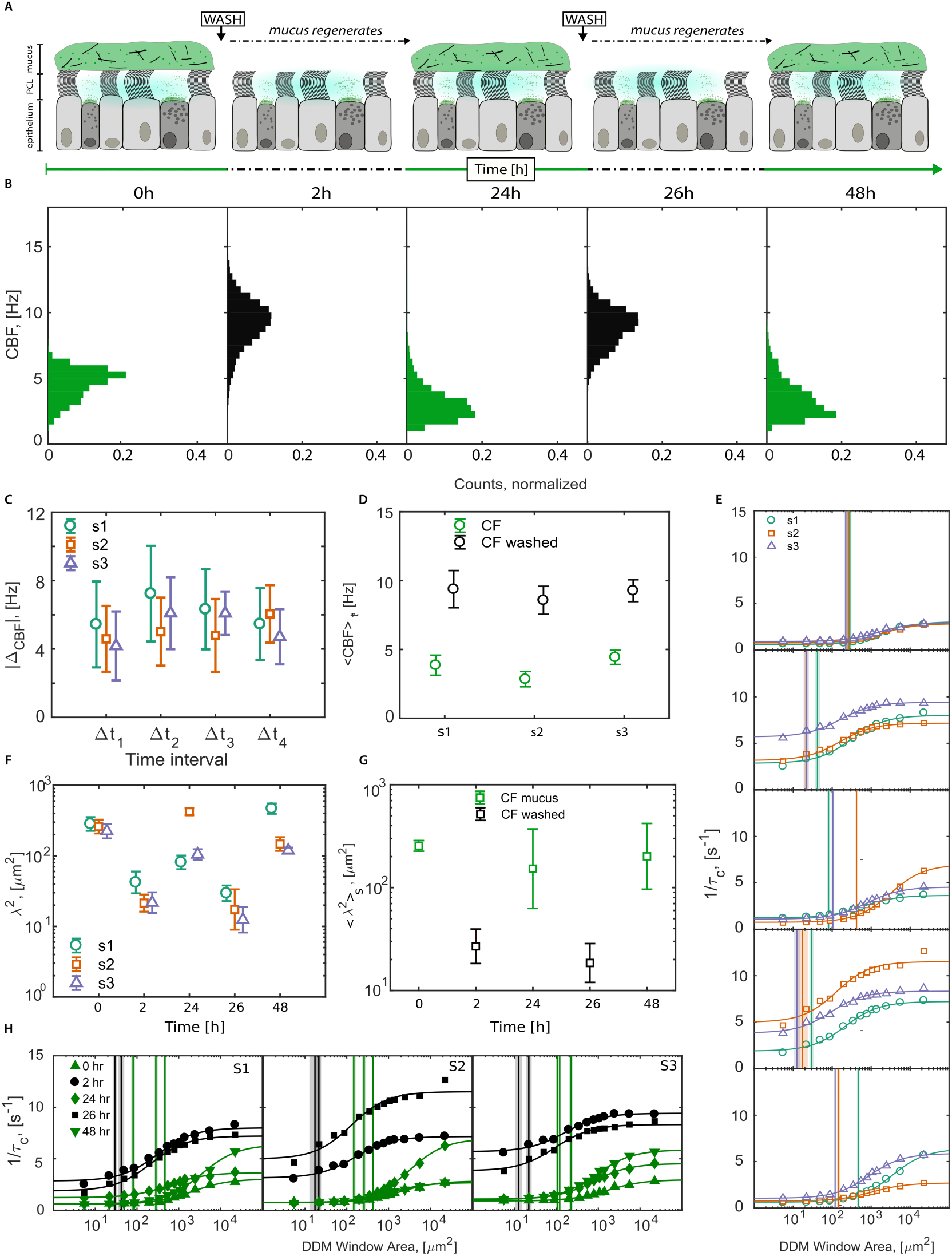
Mucus removal increases CBF and decreases the length scale of ciliary coordination. The experimental timeline of the wash/regeneration experiments is shown in **(A)**. Samples obtained from three subjects affected by CF (F508del/F508del) were subjected to successive rounds of wash and regeneration and imaged at 0, 2, 24, 26, and 48 hours (h). (**B)** The distribution of CBF of all subjects evolved over the different time points in a cyclical manner – i.e., cycling between lower CBF in the presence of mucus (green) to a higher CBF after the mucus had been removed (black), and returning to a lower CBF once mucus had regenerated (green), and so on. Each distribution was built from at least 20 fields of view imaged across each of the three samples. **(C)** The difference in the absolute value of CBF upon mucus wash/regrowth was consistent across the entire experiment and between different subject samples (symbols) regardless of the time interval. (**D**) The weighted average (circles, whiskers mark the uncertainty) of the CBF for each subject sample (S1 – S3) across time points 0h, 24h, and 48h (green, in the presence of mucus) and across time points 2h and 26h, (black, no mucus) was significantly different between the two conditions, and showed no significant difference between sample subjects. (**E**) Sigmoidal curves of the inverse of the time constant as a function of the area of the tile DDM was applied to. Experimental data points (symbols, error bars showing 25^th^ and 75^th^ percentiles were omitted for sake of visual clarity), are fitted (continuous line) in order to find the square of the coordination length scale *λ* (vertical line, the shaded region indicates its uncertainty). *λ*^2^ consistently shifted towards smaller values after removing the mucus layer, and increased again after the mucus layer regenerated. The decreased and increased length scale dependant on mucus removal and regeneration is highlighted in (**F**), where *λ*^2^ is plotted against time separately for each subject. The unpaired two-tailed Student’s t-test, run comparing log_10_(*λ*^2^) between subsequent time points, returned P values of respectively 0.01, 0.04, 0.02, and 0.008. (**G**) The geometric mean of *λ*^2^ (squares, whiskers mark the geometric standard deviation) across subjects. (**H**) All sigmoidal curves of the same subject (S1 – S3) together with the vertical line indicating the left shoulder point are plotted in the same set of axes, highlighting the shift in the length scale of ciliary coordination dependant on the presence (green) or absence (black) or mucus.

The sigmoidal curves generated by DDM analysis are similarly cyclical in nature, and the difference in the coordination length scales is statistically significant and reproducible with successive rounds of mucus removal and across different donor samples. Analysis of the time points immediately prior to and after the removal of mucus produced sigmoidal data that were cyclical in nature and very similar across subjects (Fig. 3E). Consistent with our previous observations, the length scale over which ciliary movement is correlated decreases upon removal of the mucus and increases once mucus has regenerated. These observations are quantified by plotting the left shoulder values for each subject at the various time points (Fig. 3F, and averaged across subjects in Fig. 3G). As expected, the data from cells in the presence of mucus cluster separately from data following removal of mucus, with a statistically significant difference (P<0.05) in cilia coordination between each two consecutive time points. This analysis shows that the progressive accumulation of mucus increases the length scale over which cilia are coordinated, and that the temporal dynamics of this process can be reliably detected and quantified by multi-DDM. Robustness of the multi-DDM approach is demonstrated by finding that the sigmoidal curves within each subject sample, generated for each condition, cluster closely together (Fig. 3H) and that there is no significant difference in the left shoulder points across subjects in either washed or unwashed conditions (Fig. 3F, H).

### Multi-DDM detects partially restored ciliary beating following treatment with CFTR-modulating drugs and Thymosin-α1

Having demonstrated the usefulness of phenotyping cilia dynamics and coordination with multi-DDM, not just under static conditions but also over time and in response to experimental variables, we turn to demonstrating screening and assessment of putative therapeutic compounds for the treatment of respiratory diseases, or indeed any condition in which ciliary beating plays an important role. Recent studies have reported that the CFTR-modulating drug VX-770 together with VX-809 can alleviate pulmonary exacerbations in patients homozygous for CFTR mutation F508del *(27, 28)*, and the combination has been approved by the FDA for use in patients over the age of 12. We reasoned that treating F508del/F508del HAECs with the combination of VX-770 and VX-809 (hereafter referred to as VX-770/VX-809) would be a good test of whether multi-DDM can be used to assay drug responsiveness based on the quantitative assessment of restored ciliary beating. A single, combined treatment of VX-770 (0.1µM) and VX-809 (3µM) was administered to HAECs from three subjects with the F508del/F508del mutation and data were collected at 0, 24 and 48 hours. At 0hrs, a range of CBFs from approximately 3.9 – 5.6 Hz is detected across all samples in both conditions, however at 24 and 48 hours the HAECs treated with VX-770/VX-809 show an increase in CBF, with the greatest difference compared to control seen at 48hrs (Fig. 4A, B). On average, the CBFs of the F508del/F508del HAECs treated with VX-770/VX809 for 48 hours increase by 2.6Hz compared to control, while the shape of the distribution of the CBFs for both conditions do not change markedly, suggesting that the increase in CBF occurs across the entire population. Sigmoidal curves show changes in ciliary coordination upon treatment with VX-770/VX-809 at 48 hours for each of the three different subjects (Fig. 4C). The curves representing data from three different subjects show a shift to smaller coordination scales for subjects 1 and 2 in response to the treatment. In contrast, subject 3 showed a very slight shift to larger coordination scale at 48h; this is a considerable improvement over the behaviour of the same, non-treated subject (Fig. 4D, E). These data are consistent with a partial restoration of normal ciliary beating as suggested by the CBF data from the same experiment (Fig. 4A, B) and as seen in the mucus wash/regeneration experiments (Fig. 3). Analysis of the average value for the shift of each curve with respect to the 0h time point across the three subjects reveals that, while the control treatment does not produce any effect after 48hrs, cells treated with VX-770/VX-809 show a clear difference in ciliary coordination which is detectable even when the data from all subjects were combined (Fig. 4E). Taken together, these data show how multi-DDM data can be used to detect changes in CBF and cilia coordination in F508del/F508del HAECs undergoing drug treatment either in a single subject or averaged across a group of subjects and thus how multi-DDM provides a quantitative read-out of drug efficacy.

**Figure 4.**
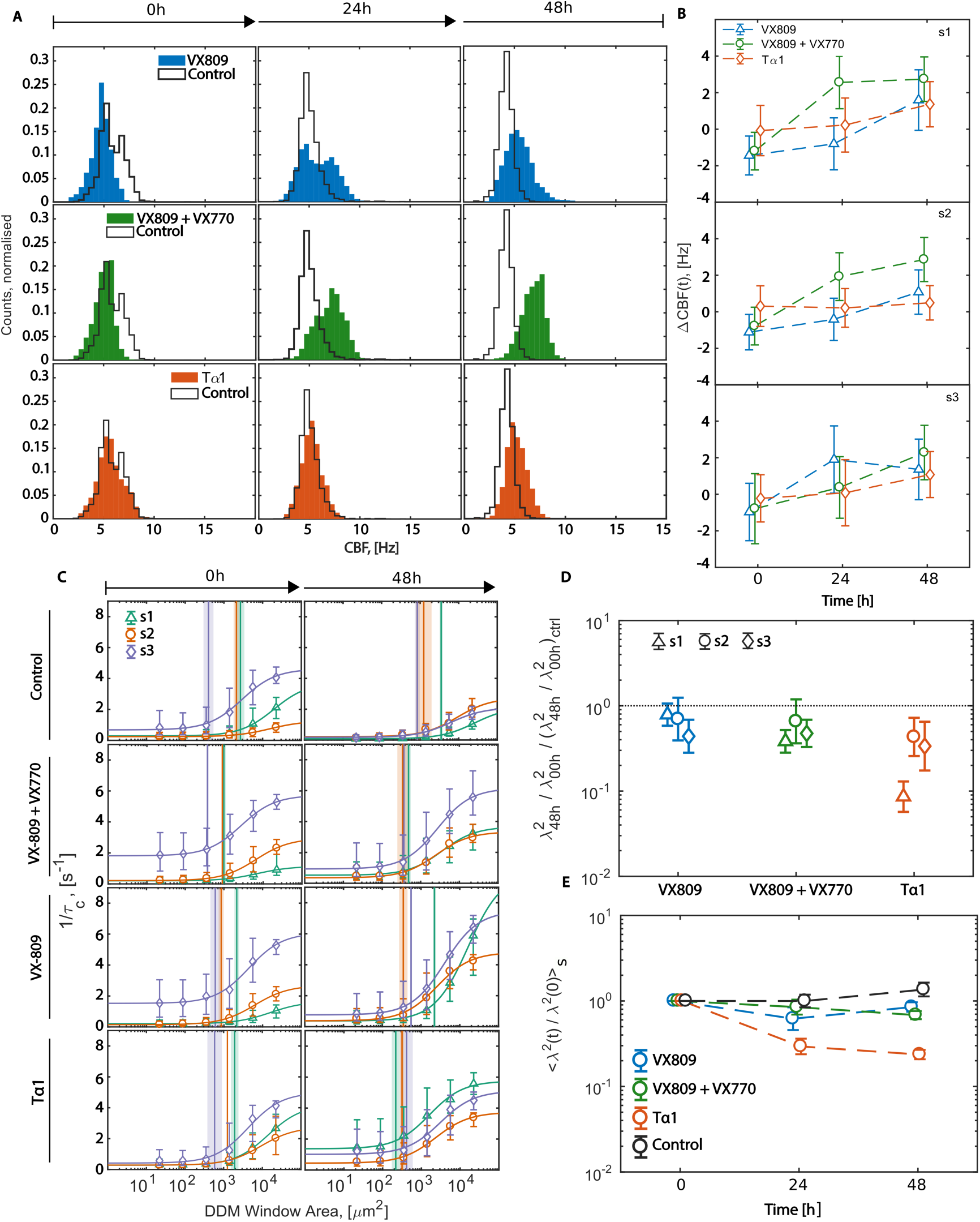
The effect of CFTR-modulating drugs on ciliary beat dynamics as assessed by multi-DDM. (**A**) The time evolution at 0, 24 and 48 hours of the CBF distribution across samples obtained from subjects with CF (F508del/F508del) treated with different CFTR-modulating drugs (VX-809, VX-770/VX-809, and Tα1, coloured plots) compared to control (DMSO, black outline). Each distribution is built from at least 16 FOVs imaged across each of the three samples. All treated samples are characterised by higher CBF with respect to the control at 24 and 48 hours. (**B**) The difference in CBF between each treated sample and the control sample (ΔCBF) increases over time for each subject sample (symbols). **(C)** Sigmoidal curves of the inverse of the DDM time constant as a function of the area of the analysed tile showed how treatment with VX809, VX809 and VX770, and Tα1 decreases the length scale of ciliary coordination, similar to what was observed upon removal of mucus (see Figure 2). (**D**) Plot shows the ratio between *λ*^2^ measured at 48h and 0h for each subject and treatment separately, normalised by the same *λ*^2^ ratio measured on the control. All data points are below 1 (dashed grey line). For each treatment, every subject showed an improvement in ciliary coordination relative to the DMSO-control at the same time point. **(E)** The time evolution of the length scale of ciliary coordination showed the greatest response to Tα1, as determined by the ratio between *λ*^2^(t) and the initial *λ*^2^(t = 0h) geometrically averaged over the three donors (whiskers mark the geometric SD) against time.

VX-809 has previously been shown to increase the amount of F508del-CFTR protein at the cell surface in HAECs with the F508del-CFTR mutations *(29)*, however a later clinical study showed that treatment with VX-809 alone is insufficient to produce any benefit in patients with the F508del mutation *(30)*. In the study, the authors reported no significant increase in CFTR function in the nasal epithelium, however interestingly CFTR function was improved in the sweat gland. Intrigued by this result, we tested the effect of VX-809 alone on cilia beating to determine whether we could detect any changes. An Ussing chamber analysis of VX-809 on HAECs with F508del/F508del demonstrates the expected properties, as observed by changes in short-circuit current in response to acute treatments with a CFTR activation cocktail (forskolin/IBMX), a CFTR potentiator (VX-770), and a CFTR inhibitor [CFTR(inh)-172] Supplementary Fig. 4A). Low baseline levels of functional CFTR expression are observed, which increase after treatment with VX-809 for 24 hours (Supplementary Fig. 4B). These results demonstrate a response to VX-809 in the primary nasal epithelial cells used for these studies, as indicated by an increase in CFTR activity. Consistent with this, multi-DDM analysis reveals changes in CBF and cilia coordination in response to treatment with VX-809 for 24 and 48 hours (Fig. 4). CBF increases slightly to an average of 5.5Hz at 48 hours, compared to vehicle-only control CBF of 4.2Hz (Fig. 4A, B). Analysis of the sigmoidal curves reveals a shift to smaller length-scales for subjects 2 and 3 upon treatment with VX-809 (Fig. 4C - E). In contrast, subject 1 shows a slight shift to larger length-scales, albeit less pronounced than the shift of the same subject under control conditions. The behaviour of the coordination length scale with time in VX-809-treated cells differs significantly from vehicle-only controls, as indicated by different trends observed when plotting the ratio between λ^2^ and λ^2^ at 0 hr, averaged across subjects (Fig. 4E). Although the effect at 48 hours is not as great as in the cells treated with VX-770/VX-809, these data nonetheless indicate that cilia coordination is altered following 48 hours of continuous VX-809 treatment in HAECs obtained from subjects with the F508del/F508del mutation in CFTR.

Thymosin-*α−*1 (Tα1; ZADAXIN) is a naturally occurring polypeptide that is used as an immunomodulator in viral infections, immunodeficiency and malignancies *(31, 32)*. Recently, Tα1 was shown to rectify the functional defects of F508del-CFTR in HAECs by increasing the half-life and the activity of the F508del-CFTR protein *(19)*, however whether Tα1 treatment affected ciliary beating dynamics was not assessed. We treated HAECs obtained from three different subjects homozygous for the F508del/F508del mutation in CFTR with 100ng/mL Tα1 over a period of 48 hrs and compared this with the same cells treated with a vehicle-only control. The range of average CBFs for subjects is 4.5 – 5.8 Hz at 48 hours following Tα1 treatment, compared to 4.1 – 4.4 Hz for the vehicle-only control (Fig. 4A, B). Analysis of the sigmoidal curves reveals a shift to smaller length-scale for coordination in all subjects, but most notably in subject 1 upon treatment with Tα1 (Fig. 4C - E). Consistent with this, the average difference between the left shoulder point before and after the treatment differs significantly from the control at both 24 and 48 hours, and is greater than either VX-770/VX-809 or VX-809 alone (Fig. 4E). The markedly decreased coordination length scale observed in response to all three drug treatments is indicative of partial restoration of normal ciliary beating dynamics, which is consistent with the previously reported roles of these drugs in modulating CFTR functionality in CF cells *(26, 32-34)*.

## Discussion

Probing the collective cilia beating dynamics and mucociliary clearance across cells at ALI with multi-DDM provides a direct quantitative phenotyping of cilia function and represents a new approach to determine the coordination of cilia and their efficiency in propelling mucus. Our framework is not limited to the estimation of the CBF in ALI culture, but also gives information about the coordination and the spatio-temporal correlation of the ciliary beating as well. This is demonstrated by the analysis of cilia coordination either in the presence or absence of mucus. In cells derived from subjects with CF, the very thick and stiff mucus increases the length scale over which ciliary movement is correlated.

In this study, we encountered two instances where our analysis was limited by extremely thick mucus: firstly, at the perimeter of the well insert in ALI culture, where mucus accumulates more rapidly than on the rest of the insert membrane. This was easily overcome by avoiding this area for data collection. The second instance in which thick mucus was limiting was in the context of CF, where cells accumulated excessive amounts of mucus when left without intervention for several days. We observed this for example in our vehicle-only controls for the drug-treatment experiments. After 48 hours, motion in the FOV for vehicle-only HAECs showed a very high lengthscale of coordinated ciliary beating, which made it impossible to use the right shoulder as it fell out of the detection range. Instead, we used the left shoulder, which represents the length scale at which perfect coordination begins to be lost. Focussing on the relative difference between shoulder points, and not the absolute values, affords greater flexibility across a range of different conditions as either the left or right shoulder point can be chosen, depending on the specific circumstances.

Mutations in CFTR are typically divided into different classes based on the particular process they affect *(33)*. The F508del-CFTR mutation affects both protein folding and channel gating, resulting in a protein which not only does not function properly but where the majority is degraded before it can reach the plasma membrane. Given the two distinct defects caused by F508del-CFTR, it is generally believed that two complementary strategies are required to target it. The combination of VX-809 and VX-770 has been proven to be effective in addressing this requirement: VX-809 is a CFTR “corrector”, addressing (to some extent) the protein processing defect, resulting in a greater amount of F508del-CFTR protein at the plasma membrane. VX-770 is a CFTR-potentiator that improves the gating properties of the CFTR protein – not only in F508del-CFTR but in other CFTR mutations too *(29)*. Our results show the clear benefit of the combination of VX-809 and VX-770, with increased CBF and cilia coordination length scale closer to healthy conditions becoming apparent after only 24 hours of treatment. An even greater effect was observed following treatment with Tα1. Unlike in our experiments on the removal and regeneration of mucus, we did not observe the total restoration of ciliary beating upon treatment with any of the three different drugs. We attribute this to the fact that, either singly or in combination, VX-809 and VX-770 can restore only a proportion of F508del-CFTR protein functionality, and CFTR activity is never rescued completely to the level seen in cells from healthy subjects. The same is true for Tα1 (26). There has been some interest in the potential effect of VX-809 alone *(29)* however to date it has not been shown to result in any clinical benefits *(30)*. Our results show a modest improvement in CBF, and reduction of the cilia coordination length scale towards heathy phenotype upon treatment with VX-809. Interestingly, we observed some variation in the response to VX-809 among the HAECs isolated from the three different subjects (HAECs from subject 1 responded less favourably than the other two) whereas we did not observe such variation among VX-770/VX-809 treated cells. Understanding why certain subjects respond differently to VX-809 may help to uncover genetic factors that contribute to the range of phenotypes and pathologies exhibited by patients with CF, leading to a more personalized approach for therapeutic intervention.

The ability to quickly and efficiently determine whether a given treatment has a positive therapeutic benefit on cells with different classes of CFTR mutations is of great value. There are nearly 2000 different mutations in CFTR, and while not all mutations cause CF, there are still many CF-causing mutations that are rare, uncharacterized, and have not been tested for their responsiveness to clinically available CF drugs. Recently, Dekkers *et al*. reported an organoid-based screen to assess the efficacy of CFTR-modulating drugs and showed its effectiveness in predicting drug responsiveness for subjects with rare, uncharacterized CFTR genotypes *(18)*. The study is an excellent example of how patients that fall into this category could benefit from such a screen, however the methodology of the assay, which involves isolation of colonic crypts, cell selection and expansion, organoid culture and seeding in Matrigel droplets, is costly, requires considerable expertise and is open to variation due to the use of Matrigel and other undefined animal-based products. For these reasons, we believe the assay is unlikely to be widely adopted by practising clinicians. In contrast, our assay is based on primary HAECs grown in ALI culture, which is a simpler cell culture already widely used amongst clinicians. Furthermore, and unlike current approaches for measuring ciliary beating parameters, our approach is unbiased, automated and considers not only CBF but also the spatio-temporal dynamics of cilia coordination across the entire FOV, the latter of which conveys important information regarding collective cilia beating and fluid transport in a clinically-relevant setting. The analysis of multiple cilia beating parameters, a “deep phenotyping” of cilia dynamics, is critical in order gain a more complete picture of how different candidate drug treatments affect mucociliary clearance. This was clearly the case with regard to Tα1 treatment, where despite only a modest improvement in CBF that was not statistically significant, we observed a marked improvement in cilia coordination, even greater than that seen in response to VX-770/VX-809.

We envision a number of applications for the multi-DDM approach presented here. Firstly, it should now be possible to quantitatively assess a wide range of putative drugs for the treatment of CF based on changes in ciliary beating and coordination. *In vivo*, MCC is mediated by the coordinated action of thousands of beating cilia, and so a quantitative approach to measure ciliary beating dynamics is highly suited to the analysis of therapeutic modulators of MCC. This is particularly relevant for drugs such as ENaC inhibitors, mucus hydrators, and DNases/mucus disruptors, which do not modulate CFTR function, and so cannot be tested using Ussing chamber analysis nor the Forskolin-induced swelling assay *(34)*. A second application of our approach is in the analysis of CFTR-modulating drugs, where it may be useful as a complementary assay alongside Ussing chamber analysis. Theoretically, restoration of CFTR ion channel function as measured by Ussing chamber analysis should lead to mucus rehydration and increased MCC. In practise, however, the degree to which CFTR activity is restored varies among patients, and the relationship between the Ussing chamber results and the observed clinical response is not always clear. In the future, it would be interesting to investigate whether there is any relationship between data obtained via Ussing chamber, multi-DDM and *in vivo* efficacy to see if there is a threshold of ion transport that is required in order to improve MCC. Application of multi-DDM to assess putative drug treatments would require the culture of HAECs isolated from the subject and application of multi-DDM analysis before and after the treatment, which is quite straightforward. An additional application of multi-DDM is to investigate ciliary beating dynamics in diseases other than CF, such as in ciliopathies, where over 187 genes have been associated with a defect in cilia form or function *(1, 35)*. Characterising ciliary beating in different classes of ciliopathies and in the range of different tissues in which ciliary beating is affected will help to better understand this broad class of disease. This in turn may lead to better diagnostic tools, particularly for diseases in which the underlying genetic mutation remains completely uncharacterised.

In conclusion, we have developed a straightforward, quantitative assay based on multi-DDM to characterise and detect changes in ciliary beating of HAECs grown in ALI culture in both healthy and disease contexts. Our approach improves on previous methods of ciliary beat analysis in that it is unbiased and automated, and because it captures the spatio-temporal coordination of collective cilia beating which is crucial for MCC, and not simply the CBF. We provide a proof-of-principle by applying multi-DDM to HAECs obtained from subjects with the F508del/F508del mutation in CFTR to assess their responsiveness to known CFTR-modulators VX-770, VX-809 and Tα1. Our data indicate that multi-DDM can quickly and efficiently detect compounds that result in restored ciliary beating dynamics – in this case VX-770/VX-809 and Tα1, and to a lesser extent VX-809 alone. Our approach provides a straightforward, quantitative assay to assess patient responses to putative and existing drug treatments for CF, which may be applied to CFTR-modulating and non-CFTR-modulating drugs alike. Finally, since multi-DDM is ultimately a means to assess cilia beating dynamics, its use is not limited to CF but may be applied to other diseases in which ciliary beating is affected.

## Material and Methods

### Study design

The overall objective of this study was to use the recently-developed multi-DDM algorithm to characterise ciliary beating in cells from subjects with CF, and to test whether the approach was sensitive enough to detect changes in ciliary beating over time and in response to various drug treatments. As a first-pass, we applied multi-DDM to primary HAECs purchased from Epithelix and manipulated the mucosal layer to affect ciliary beating. Once our approach was validated – i.e., it became clear that the data obtained from multi-DDM analysis could be used to distinguish between different samples under different conditions - we obtained primary clinical samples for experiments involving CFTR-modulating drug treatments. For every experiment, the DDM analysis was based on data collected from HAECs obtained from three different subjects, either healthy subject controls or cells harbouring the F508del/F508del mutation in CFTR. We chose to focus on this mutation as it is the most common cause of CF and one for which at least one therapeutic drugs has been approved (ORKAMBI^®^). At least sixteen - often more - videos were taken of each of the samples at each data collection point, avoiding the perimeter of the cell culture insert due to the accumulation of mucus in that region (an experimental artefact caused by the geometry of the vessel). Multi-DDM analyses were performed as previously described *(15)* and in a such a manner as to ensure the entire field-of-view was represented in an unbiased manner. This study was not blinded.

### Human material and cell culture

HAECs obtained through the clinic were isolated from nasal brushings of the inferior turbinate performed using a cytology brush on CF subjects attending the Adult Clinic at National Jewish Health (Denver, CO, USA). Samples were collected using a protocol approved by the Institutional Review Board at National Jewish Health, and all subjects provided written informed consent. The median age of the donors was 41 years old, 4/6 were female, and all were Caucasian and of the CFTR genotype F508del/F508del. Nasal epithelial cells obtained from brushings were expanded in culture using previously described methods of conditional reprogramming culture (CRC) *(36)*. Briefly, nasal brushing samples were dissociated in Dulbecco’s phosphate buffered saline solution containing 5 mM EDTA and 5 mM EGTA and mechanically processed by repeatedly pipetting. After washing, cells were resuspended in CRC medium containing the rho-kinase inhibitor, Y-27632, and plated on a feeder cell layer of gamma-irradiated 3T3 fibroblasts. Reprogrammed epithelial cells were expanded at 37 °C in a humidified incubator in an atmosphere containing 5% CO_2_, removed using two-stage trypsinization, and replated on a fresh feeder layer. After reaching ∼80% confluence, passage 2 cells were aliquoted and stored in liquid nitrogen. Later, thawed cells were plated onto feeder layers in CRC media with Y-27632 and expanded until ∼80% confluence was reached. After a two-stage trypsinization and treatment with DNase I, cells were plated onto collagen-coated permeable supports in a 24-well format (Corning Incorporated, Tewksbury, MA, USA) at a density of 100,000 cells/cm^2^. Once confluence was reached, apical media was removed and basal media was switched to PneumaCult ALI media (STEMCELL Technologies). Cells formed well-differentiated cultures after 4 – 6 weeks at the air-liquid interface. HAECs obtained from Epithelix (MucilAir™, Epithelix Sàrl, Geneva, Switzerland) were isolated from bronchi and cultured as per the manufacturer’s instructions. General cell culture maintenance involved washing the cells twice a week as previously described *(37, 38)*. Briefly, the apical surfaces of the cultures were incubated for 20 min in 200μl of sterile Phosphate-Buffered Saline, PBS (GIBCO^™^). After 20 minutes, the solution was gently removed by suction pipette. In preparation for data collection, cells received an additional wash the day before the experiment: the apical surfaces of the cultures were washed twice in PBS containing 1mM Dithiothreitol (DTT) solution (Sigma Aldrich, St. Louis, MO) and two additional PBS washes were applied to remove all free DTT and remaining mucus.

### CFTR-modulating and Thymosin-α1 assays

Solutions of VX-809 (lumacaftor; Selleck Chemicals LLC, Huston, USA), VX-809 and VX-770 (ivacaftor; Selleck Chemicals LLC, Huston, USA) were prepared in DMSO (Sigma) at 3mM, 0.1mM concentrations respectively. Thymosin α1 (Tα1, CRIBI Biotechnology, Padova, Italy) was reconstructed in MilliQ water at 100µg/ml concentrations *(19)*. Each compounds were then diluted 1000-fold in culture media and added basolaterally to HAECs in ALI culture at 0, 24, and 48. The control treatment consisted of dimethyl sulfoxide (DMSO) 0.1% (Sigma). CFTR-mediated ion transport was measured using an Ussing chamber system (Physiological Instruments). Cell inserts were bathed in symmetrical Ringer’s solution (120 mM NaCl, 10 mM D-glucose, 3.3 mM KH_2_PO_4_, 0.83 mM K_2_HPO_4_, 1.2 mM CaCl_2_, 1.2 mM MgCl_2_, and 25 mM NaHCO_3_), gassed with 5% CO_2_/95% O_2_, and maintained at 37°C and pH 7.4. Inserts were voltage clamped and short-circuit currents were measured. Sodium transport through the epithelial sodium channel was inhibited using 100 µM amiloride, and after currents became stable, CFTR function was measured by activation using a cocktail of 10 µM forskolin/100 µM 3-isobutyl-1-methylxanthine, followed by potentiation using 1 µM VX-770, then inhibition using 10 µM CFTR(inh)-172.

### Video acquisition

At least 20 videos were acquired from each samples at time 0h, 24, 48 and 72h on a Nikon Eclipse Ti-E inverted microscope (Nikon Instruments, Japan) with a 40x objective (Plan Apo λ 40x, N.A. 0.95, Nikon). Digital high-speed videos were recorded under bright field illumination at a sampling frequency of 150fps using a Grasshopper^®^3 GS3-U3-23S6M-C CMOS camera (FLIR Integrated Imaging Solutions GmbH, Germany). Samples of epithelial cells were imaged in a custom made chamber, where temperature, CO_2_ and humidity were continuously monitored and maintained at values of 37°C, 5% and 90% respectively. Any videos that showed drifting or duplicated areas of analysis were not included in the analyses.

### Data processing

Videos were uploaded to our custom multi-DDM algorithm pipeline (coded in MATLAB, the MathWORKS) and processed as previously described *(15)*. Briefly, the DDM algorithm prescribes to take the algebraic difference of several couples of frames separated by a lag time τ. These differences are then Fourier transformed in space, and the results averaged, to yield an averaged 2D power spectrum. If the anisotropy of the sample’s dynamics is not of interest for the analysis, an azimuthal average of the 2D power spectrum is taken. By repeating the process for different values of the time lag *τ* we build the Image Structure Function *I(q,τ) (39, 40)*. As detailed in *(15)* we then extract information about the dynamics of the sample by fitting the *I(q,τ)* with an empirical function

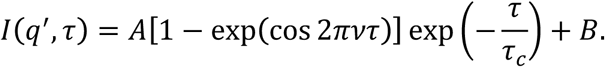

The frequency of oscillations ν is the CBF, and the time constant *τ*_*c*_ measures the degree of coordination within the field of view. The MATLAB code we developed performs the DDM algorithm automatically on the biggest square region that the fields of view can contain, and on each tile of increasingly finer square grids. The software produces an output file per each video. Each output file contains the result of the DDM algorithm for each of the tiles (of all sizes) analysed. The software also gives the user a list of the tiles that were covering a region of the sample with little or no motion, and/or where the fitting of *I(q,τ)* failed or was deemed unreliable. These tiles were not used for subsequent analysis.

The motion detection used for the experiments in Fig. 1 - 3 is described in *(15)*, and relies on thresholding an image obtained by taking the standard deviation of the fluctuation of each pixel’s recorded grey level over time. This simple approach works well in most cases, but struggled in some videos during the experiments featured in Fig. 4, because it did not discard some DDM tiles where there was no ciliary motion. We therefore devised a different motion detection algorithm, geared towards better performances in case of low signal-to-noise ratio. In this revised algorithm we take differences between every other frame, to highlight regions where the sample shows fast motion. These images undergo a median filtering, to reduce the effect of camera noise, and a local standard deviation filtering. This last filter highlights regions with high alternation of dark and bright pixels, where the moving cilia are located. We then take the base-10 logarithm of the standard deviation across all these filtered images. The motion map thus obtained is bright in correspondence of moving cilia, and dark over static regions of the sample. Segmentation then finds the foreground features, where the beating cilia are. This is done by fitting the dark end of the pixel values histogram with a Gaussian, and marking as background (so, no moving cilia) all pixels with value smaller or equal than the sum of mean and width of the fitted Gaussian. This algorithm was seen to reliably work as long as some static regions are present in the field of view. Once all videos taken on a sample were automatically analysed we collected the CBF values measured on DDM tiles (of size closely matching a cell size, so 64px, or 9.3µm side) across all videos pertaining to the same sample to build the CBF distributions.

The sigmoidal curves showing the loss of coordination upon increasing the DDM tile size are built by pooling together data from many fields of view on the same sample. For each analysed tile size, we build the distribution of values of decay time τ_c_ measured on tiles of matching size across all videos on the same sample. The marker in the plots shows the medians of such distributions, while the whiskers show the 25^th^ and 75^th^ percentiles (see an example in Fig. 1H and Fig. 4C). The experimental data points are then fitted as 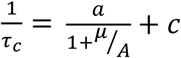, where a and c give the level of the left and right plateaus, *A* is of the area of the DDM tile, and *µ* is the inflection point of the sigmoid. The left and right shoulders of the sigmoidal curve are then the abscissas (*λ*^2^ = *μe*^−2^ and Λ^2^ = *μe*^2^) of the points where the line that best fits the central slope meets the prolongation of respectively the left and right plateaus.

### Statistical analyses

CBF results are presented as mean ± SD, unless specified otherwise in the figure caption. *λ*^2^ measurements on a single sample are presented as measurement ± 68% confidence interval, while averages across subjects are presented as geometric mean ± geometric SD. Statistical analysis were performed by paired or unpaired two-tailed Student’s t-test using MATLAB, and by weighted one-tailed t-test using R. The 95% confidence level was considered significant. Details about P values and which test was used to calculate them are in the figure captions.

## List of Supplementary Materials

**Supplementary Table 1:** CBF of samples featured in Fig. 2

**Supplementary Table 2:** CBF of samples in Fig. 3

**Supplementary Table 3**: CBF of samples in Fig. 4

**Supplementary Figure 1:** distribution of CBF measured before and after washing

**Supplementary Figure 2:** time evolution of the CBF distribution

**Supplementary Figure 3:** sample specific time evolution of the CBF distribution during drug screening

**Supplementary Figure 4:** Ussing chamber analysis for VX-809

## Acknowledgments

We thank Dr C. E. Hendry for rigorous and thorough manuscript edits

## Author contributions

M. C. and L. F. performed the experiments, interpreted the data, wrote the article text and generated figures. M. C. designed the overall study. L. F. developed the analysis software. P. E. B. provided patient material, performed USSING chamber analysis and provided the associated data. J. K. designed and optimised the high speed video microscopy (HSVM) apparatus, the HSVM acquisition software and built the microscope incubator chamber. P. C. obtained funding, interpreted the data and provided overall steering and support

## Data and materials availability

DDM software and digital videos data used in this study are available upon request.

